# Direct comparison of CRISPR knockout and interference with Perturb-seq

**DOI:** 10.64898/2026.07.04.736492

**Authors:** Laura M Drepanos, Berta Escude Velasco, Abigail J Chase, Smriti Srikanth, Michael Gatzen, Hannah D Rickner, Dan Dubinsky, Andrew W Navia, Tsukasa Shibue, Kathleen B Yates, Peter S Winter, John G Doench

## Abstract

CRISPR knockout (CRISPRko) and CRISPR interference (CRISPRi) are two workhorse technologies for loss-of-function studies, yet direct comparisons between the two are scant relative to their widespread adoption. Here, we establish benchmarking libraries for Cas9-based CRISPRko and CRISPRi screens using Perturb-seq as the read-out. For both modalities, we observe consistent transcriptional signatures among cells with the same genes perturbed, strong evidence of on-target signal. We also examine tradeoffs between modalities: while CRISPRi guides demonstrate heightened rates of off-target activity, we also observe artifacts stemming from the cellular response to double-stranded breaks with the use of CRISPRko. The libraries and analyses presented here will be a useful benchmarking and de-risking resource for any group preparing for a large-scale Perturb-seq screen.

## Introduction

CRISPR knockout (CRISPRko) and CRISPR interference (CRISPRi) are both effective approaches to introduce loss-of-function mutations^1,2^. Particularly in light of recent advances in library design, both modalities enable low false negative screening rates^3,4^. These complementary technologies involve distinct mechanisms of action and consequently offer numerous tradeoffs. The introduction of a double-stranded break (DSB) with CRISPRko establishes permanent loss-of-function through the introduction of insertions and deletions (indels) by imperfect DNA repair. A limitation of this approach is that DSBs may yield an antiproliferative effect, an especially pertinent issue when targeting genes in copy number amplified regions^5^. Moreover, CRISPRko can impact expression of genes telomeric of the target site in a fraction of cells, an artifact that has been postulated to reflect large-scale chromosome modifications^6–8^. CRISPRi, in contrast, avoids toxicity associated with DSBs by employing a deactivated Cas9 nuclease (dCas9) to actively block transcription. However, the transcription start site (TSS) of a target gene is sometimes poorly defined or heterogeneous, and genes that share promoter regions cannot be targeted individually^9,10^.

Perturb-seq experiments inform the phenotypic effect of perturbations introduced by CRISPR single guide RNAs (hereafter simply “guides”) by identifying the guide and a portion of the endogenous RNA transcripts present in each cell^11–15^. The initial cost estimate of $0.05 per cell^11^ proved overly-optimistic at the time, but subsequent methodological improvements and decreases in sequencing costs suggest that wider-spread adoption of this approach is now within reach for the wider community^15^. The high dimensional readout of Perturb-seq enables unbiased detection of perturbation responses, making it a suitable method to compare screening modalities.

Here, we report head-to-head Cas9 CRISPRko and CRISPRi Perturb-seq screens to compare the rates of on-target and off-target activity between these screening technologies. While the data presented examines a relatively small library, its deliberate design enables detailed comparison of these technologies without confounding experimental factors. Moreover, we leverage the use of four constructs per target gene to define guide-specific off-target effects as those not shared with others targeting the same gene. We present longitudinal data up to fourteen days post-transduction to broadly inform the screening behavior of either modality.

## Results

### Library design and validation

We set out to develop nimble but informative benchmarking libraries that could be used across a wide spectrum of human cell models to qualify the performance of CRISPRko or CRISPRi. To facilitate an informative comparison of on-target activity, we chose to target genes with prior evidence of large transcriptomic effects in the K562 leukemia cell line^10^. From this set, we identified 25 genes that represent a broad and largely non-overlapping range of known functions (Figure 1a). This gene set features 23 nonessential genes to enable profiling at late time points, as well as two essential genes (*RPL9, DCAF13*) as positive controls for a strong transcriptional effect prior to their depletion^16^ (Figure 1b). We selected four guides per gene per modality, notably incorporating some, but not all, of the advances we have recently reported for CRISPRko and CRISPRi guide design^3,4^, as this work was conducted concurrently. The libraries additionally employ ten negative control guides targeting intergenic sites, yielding a total of 110 perturbations in each. The guides were cloned into a Fragmid-assembled^17^ CROPseq vector architecture^18^ with a puromycin resistance cassette.

**Figure 1:**
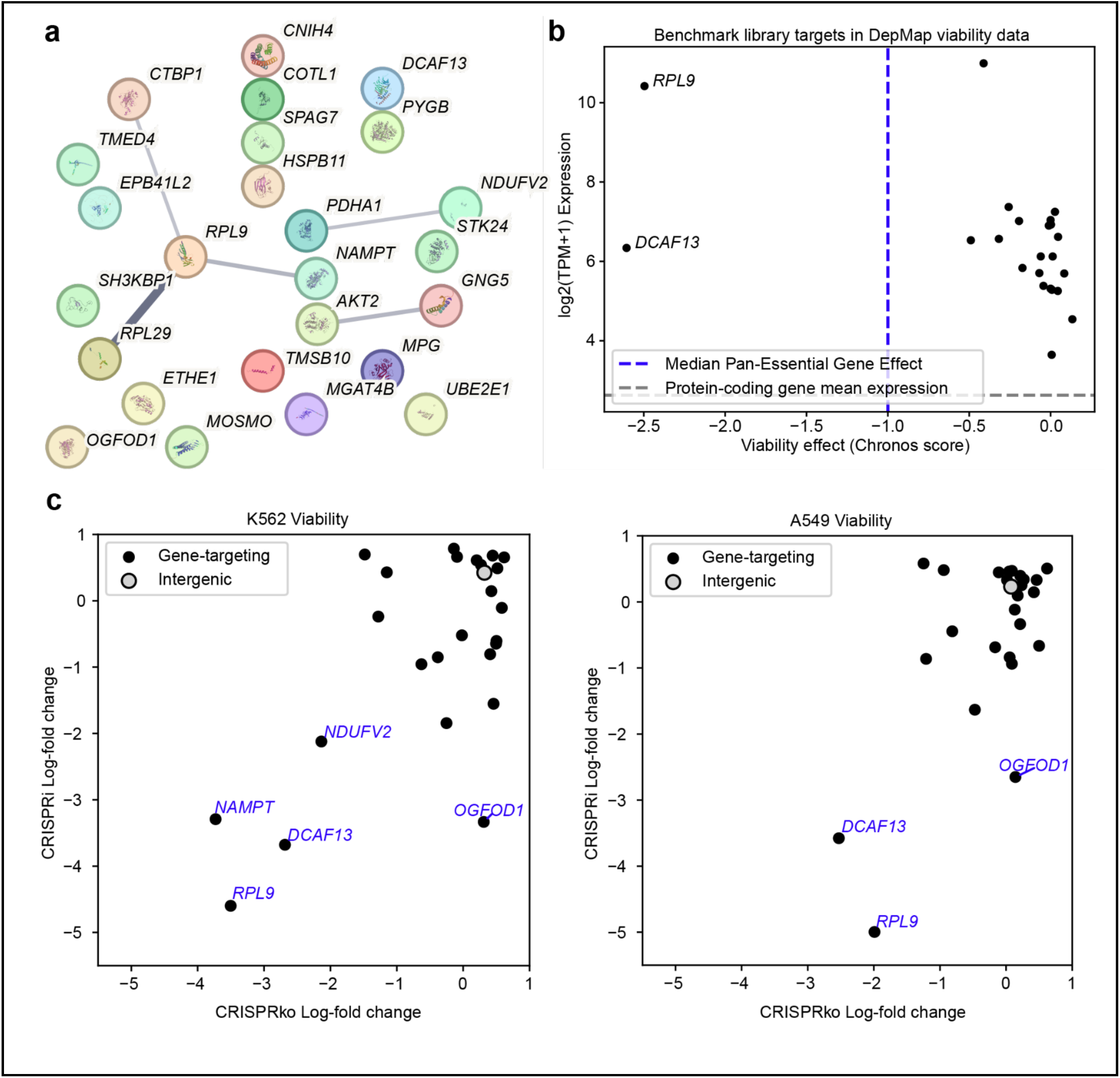
Design and validation of CRISPRko and CRISPRi benchmark libraries. a) Protein relationship network analysis of all 25 benchmark library targets. The map is generated with STRING Version 12.0 using all default interaction sources. Line thickness indicates the strength of supporting data, and thus confidence, that two genes are associated. All connections with at least medium confidence (STRING interaction score=0.400) are shown. b) Behavior of library targets in existing screening and expression data (DepMap 2026 Q1). c) Log-fold change in guide abundance relative to pDNA for all guides in the benchmark library at Day 14. Each point represents the average log fold change among guides targeting the same gene. Each plot contrasts the viability effects of perturbing each gene with either CRISPRko or CRISPRi in the indicated cell line.

Prior to initiating Perturb-seq screening, we confirmed the utility of our vectors and guides with viability screens in the K562 and A549 cell lines, as representative suspension and adherent models. The lines were engineered to express Cas9 or KRAB-dCas9, utilizing an N’terminal fusion of the ZIM3 KRAB domain for the latter^19^. In both models, we observed depletion of guides targeting the positive control essential genes *RPL9* and *DCAF13*, indicating that we successfully engineered vectors and designed libraries capable of introducing loss-of-function in this screening context (Figure 1c). We additionally noted depletion of guides targeting *NAMPT* in K562 but not A549 cells.

In both cell lines, we observed depletion of CRISPRi guides targeting the nonessential gene *OGFOD1*. A likely explanation is that the TSS of *OGFOD1* is 222 base pairs away from that of the essential gene *NUDT21*. Since CRISPRi guides that target near the TSS of a gene effectively block transcription, it is unsurprising that guides targeting *OGFOD1* additionally repress this neighboring gene^2,9^ (Figure 1d). In contrast, CRISPRko guides targeting the exons of *OGFOD1*, which do not overlap with the *NUDT21* protein coding region, did not impact viability.

### Perturb-seq screens are of satisfactory quality

We transduced K562 cells expressing Cas9 or KRAB-dCas9 with the benchmark CRISPRko and CRISPRi libraries, respectively, at low multiplicity of infection. To minimize variability during library preparation between the samples, we staggered transductions across four separate time points and harvested the cells for single cell transcriptomic sequencing on the same day (Figure 2a). As the gene-targeting guides in each library are distinct, we encapsulated and sequenced the CRISPRko and CRISPRi libraries together in two technical replicates. We opted for this atypical experimental design in order to observe any potential technical noise in the downstream cell processing steps. As an exception, at Day 14 the CRISPRko and CRISPRi experiments were encapsulated for sequencing separately to distinguish intergenic-targeting guides, which were the same in each library, to enable analysis of guide effects calculated relative to negative controls for each modality (Supplementary Note 1). We repeated this experiment in the A549 cell line with a single Day 7 time point to inform the generalizability of the observed results.

**Figure 2:**
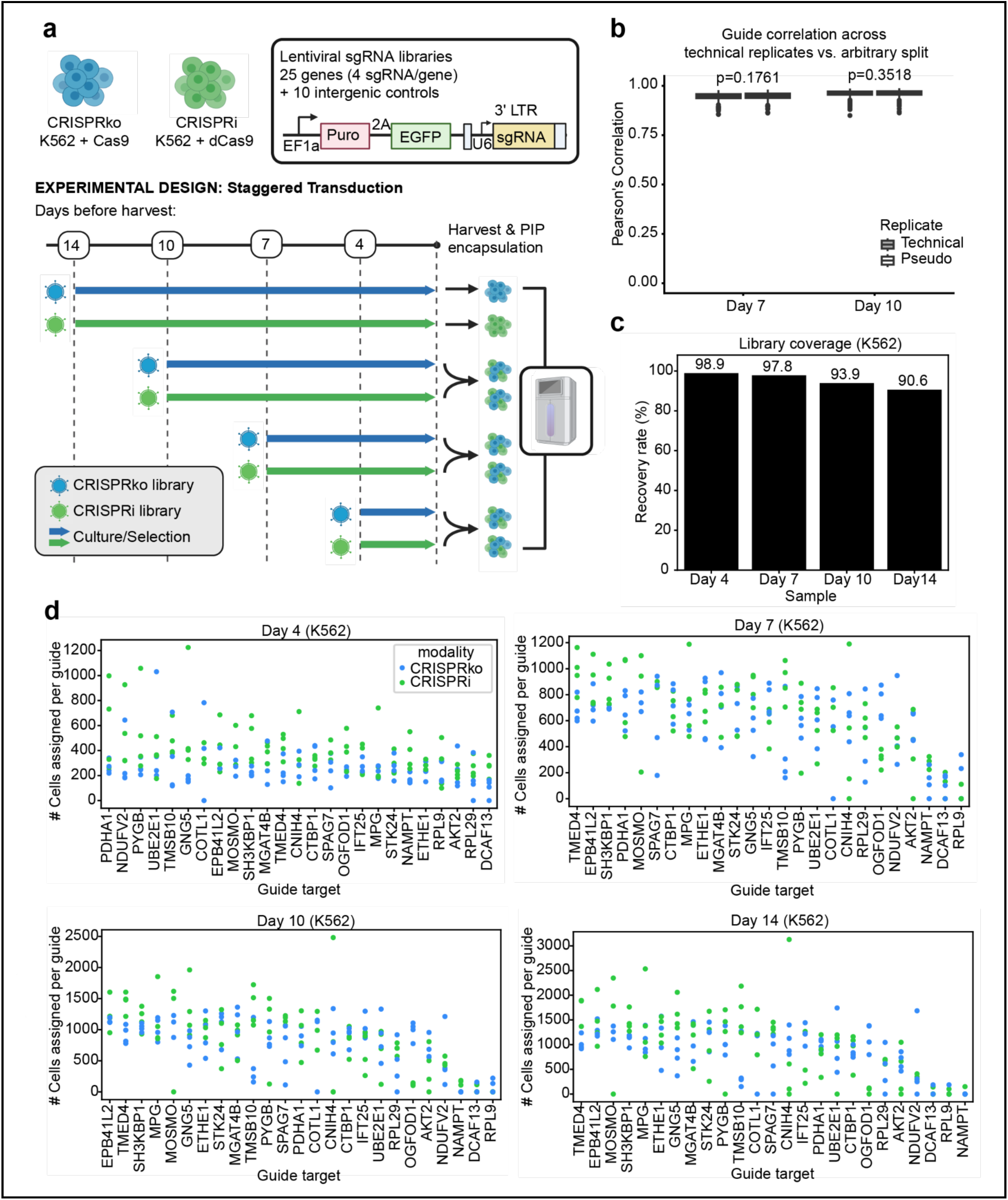
Perturb-seq data yields satisfactory quality and library coverage. a) K562 timecourse PIPseq screen experimental set up. This same design is repeated for the single-timepoint A549 experiment. b) Assessment of technical replicate correlation among samples for which replicates are joined for subsequent analysis. Using all cells that pass guide assignment from the indicated sample, cells are pseudobulked by their assigned guide and either their technical replicate or a random label (which determines the “pseudoreplicate” that the cell belongs to). Pearson’s correlation is calculated between the pseudobulked profile of cells carrying the same guide across groups on the 2000 most highly variable genes (Seurat). The correlation for each guide is shown in the plot. The correlations obtained by calculating across either technical replicate or pseudoreplicate are compared with a two-sided Mann-Whitney test. c) Percentage of guides in the benchmark library with a coverage of at least 100 cells in each sample. Samples prior to Day 14 represent the data from Replicate 1 and Replicate 2 combined, and percentages are calculated with the complete set of CRISPRko and CRISPRi guides as the denominator. Guides targeting positive essential controls (*RPL9, DCAF13*) are excluded from the calculation. d) Number of cells assigned for each guide in the benchmark library per sample. The Day 7 and Day 10 timepoints represent the total cell count from both technical replicates combined. Guides are grouped horizontally by their target gene, which are in descending order by mean coverage, and shaded according to their modality.

Cells were encapsulated using PIPseq technology^20^, which relies on direct capture of the guide RNA to generate separate gene expression and guide libraries that share a per-cell barcode. We observed adequate sequencing depth among various performance metrics across samples (Table 1). Technical replicate 1 at Day 4 in the K562 cell line exhibited disproportionately low detection of transcriptomic reads; while prior studies have reported that 1000 UMIs per cell is adequate to identify whether or not a perturbation has a substantial transcriptomic effect^21^, here we strived for higher sensitivity to detect the extent of on-target and off-target guide effects. In this instance, we reasoned that the median sequence depth of 487 genes per cell was inadequate to cover the complexity of perturbation effects since prior works have identified that loss-of-function perturbations can affect over 500 genes^16^. We therefore excluded this sample from the remainder of the analysis, leaving a single technical replicate for the K562 Day 4 timepoint. Otherwise, we observed that cells carrying the same guide across technical replicates were as similar in transcriptional space to those within the same technical replicate (Figure 2b). We therefore combined technical replicates for the remainder of the analysis.

**Table 1:** Metrics describing the sequencing depth and quality of each Perturb-seq sample employed in this experiment. Passing cells refer to droplets whose read counts resemble that of a complete cell as identified by the DRAGEN software. The assignment rate informs the fraction of cells upon removal of those with over 15% mitochondrial transcripts that are assigned a single guide. See Methods for usage of the DRAGEN software and the guide assignment protocol.

| Sample | # aligned, passing cells from DRAGEN | median genes per passing cell | median transcriptomic UMIs per passing cell | median guide reads per passing cell | # cells after removing those with >15% mitochondrial transcripts | # cells after assignment | assignment rate |
| --- | --- | --- | --- | --- | --- | --- | --- |
| K562 Day 4 Rep1 | 114001 | 487 | 1079 | 329 | [dropped] | [dropped] | [dropped] |
| K562 Day 4 Rep2 | 102875 | 1911 | 8213 | 141 | 101487 | 73772 | 72.69% |
| K562 Day 7 Rep1 | 75342 | 1880 | 6220 | 976 |  |  |  |
| K562 Day 7 Rep2 | 105761 | 1260 | 4138 | 469 | 179433 | 136285 | 75.95% |
| K562 Day 10 Rep1 | 104320 | 1597 | 4330 | 4940 |  |  |  |
| K562 Day 10 Rep2 | 101264 | 2623 | 11517 | 1771 | 203620 | 182730 | 89.74% |
| K562 Day 14 CRISPRko | 115379 | 2033 | 6311 | 899 | 114435 | 97458 | 85.16% |
| K562 Day 14 CRISPRi | 127143 | 2451 | 10929 | 847 | 126246 | 111899 | 88.64% |
| A549 Day 7 Rep1 | 130114 | 2768 | 10059 | 1245 |  |  |  |
| A549 Day 7 Rep2 | 109236 | 2822 | 11272 | 1549 | 260311 | 234346 | 90.02% |

The resulting data yielded satisfactory coverage as measured by the fraction of nonessential-targeting guides confidently assigned to at least one hundred cells per sample (Figure 2c). As expected, we observed lower recovery of cells that received guides targeting essential genes, particularly at later timepoints, suggesting successful on-target activity (Figure 2d).

### Perturb-seq screens demonstrate consistent guide on-target activity

We next determined the rate of target transcript knockdown and observed that the target transcript was among the most downregulated transcripts for most guides (Figure 3a). This knockdown was significant for a majority of gene targets (FDR <0.1, SCEPTRE^22^), particularly after Day 4 (Figure 3b). However, assessing target transcript knockdown alone likely underestimates perturbation efficacy. First, the target gene transcript may be poorly detected in unperturbed cells, thereby restricting the ability to observe significant downregulation. For example, we observed low detection of *SH3KBP1* in cells with intergenic guides, and consequently minimal target knockdown in cells with *SH3KBP1*-targeting guides (Figure 3b, Supplementary Figure 1a). Second, CRISPRko does not necessarily decrease target transcript abundance: while indels prevent expression of the full-length protein, the degree of mRNA loss is dependent on rates of nonsense-mediated decay, which varies substantially across transcripts and cell types^23^. Indeed, in some instances, feedback loops lead cells to overexpress transcripts for a gene perturbed with CRISPRko to compensate for the loss of function^24^, although we did not observe any target upregulation at FDR <0.1 in this experiment.

**Figure 3.**
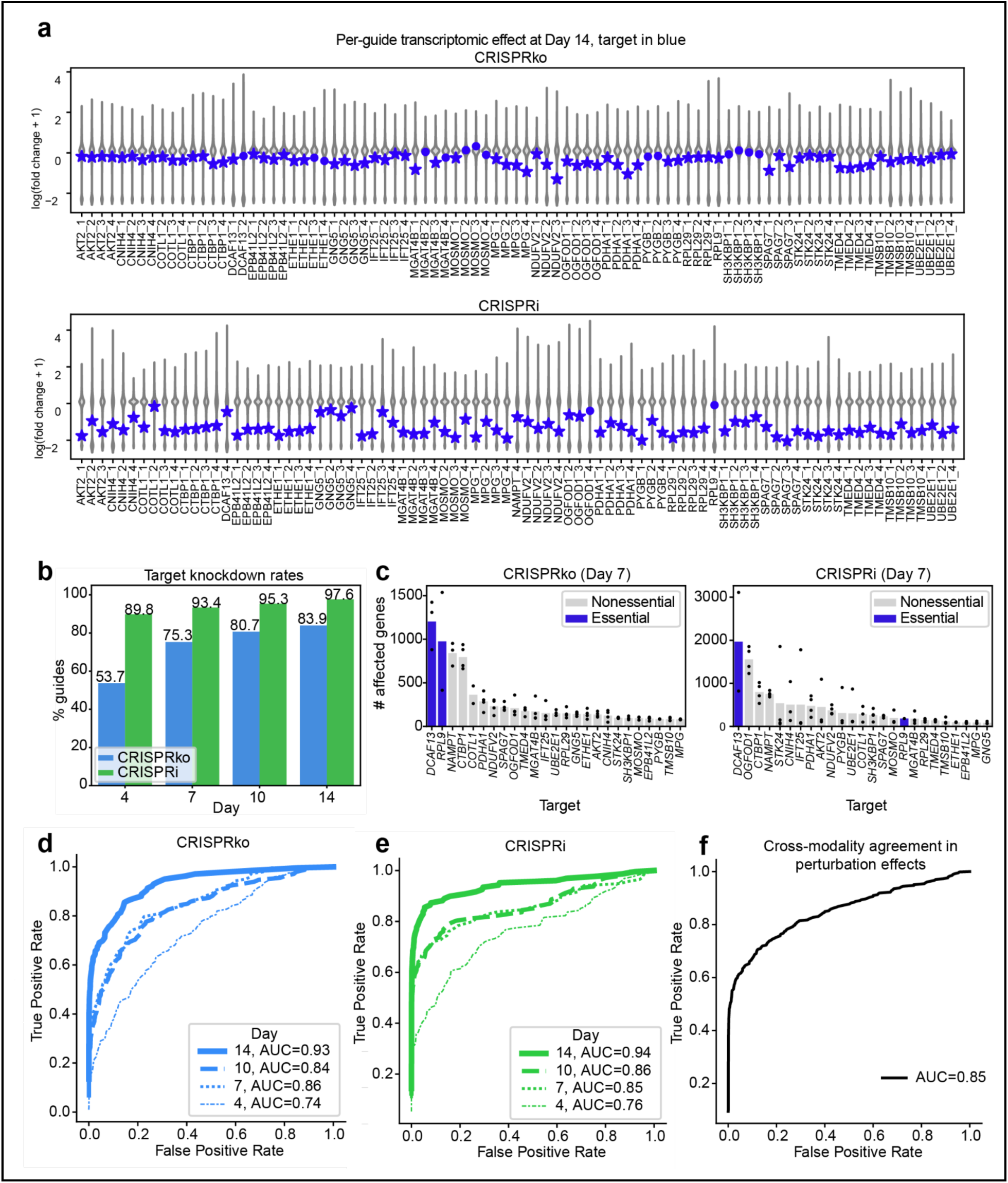
Perturb-seq reveals successful on-target activity of CRISPRko and CRISPRi guides. a) Per-guide effect on all transcripts that pass SCEPTRE pairwise quality control at Day 14 in K562 cells. SCEPTRE differential expression analysis is run in singleton mode with cells assigned intergenic-targeting guides as the reference. Log fold change therefore represents transcript abundance in cells with the indicated guide relative to cells with intergenic-targeting guides. Target transcripts are shown in blue, and transcripts with a false discovery rate (FDR) below 0.1 are denoted with a star marker. Guides not shown lack the minimum coverage of 100 cells in the indicated sample. b) The percentage of guides targeting nonessential genes that yield knockdown (SCEPTRE FDR<0.1) of the target transcript in K562 samples. c) Count of genes differentially expressed (SCEPTRE FDR <0.1) per library target. K562 Day 4 and Day 7 data is shown since essential-targeting guides deplete afterwards. Each point represents the number of differentially expressed genes from each guide targeting the indicated gene, and the height of the bar represents the average. Target genes are sorted in descending order by the average count of differentially expressed genes per guide. d) Receiver operating characteristic curve of correlation in signed p-values (as determined by SCEPTRE) for CRISPRko guides targeting the same vs. different genes. Correlation is calculated upon all transcripts that pass SCEPTRE pairwise crosswise correlation for all contrasted conditions. The results represent data from the K562 cell line. e) d, but for CRISPRi. f) Receiver operating characteristic curve of correlation in signed p-values (as determined by SCEPTRE) for CRISPRko guides against CRISPRi guides targeting the same vs. different genes. The results represent data from the K562 cell line at Day 14 to account for modality-dependent transcriptional signatures (Supplementary Note 1).

Given these limitations of assessing guide performance by examination of only the target gene, it is preferable to leverage the whole transcriptomic readout to characterize perturbation efficacy. Traditional single-cell “tissue atlas” studies typically focus on characterizing distinct cell types that have diverged functionally over millions of years of multicellular evolution, as well as lengthy developmental processes within an individual organism. In contrast, Perturb-seq experiments involve assessing the impact of a single gene alteration in a population of cells after several days. Since individual gene perturbations generally produce relatively sparse changes across the transcriptome, cells sharing the same perturbation do not drive distinct clustering using unsupervised methods like UMAP (Supplementary Figure 1b). Reassuringly, we observed that cells assigned to essential-targeting guides elicited a widespread transcriptional response prior to their depletion, a result consistent with previous studies^16^ (Figure 3c).

Beyond the magnitude of change, it is challenging to evaluate the accuracy of transcriptional responses to perturbation since objective ground truth is often lacking. We therefore assessed transcriptional similarity across guides targeting the same gene as evidence of successful perturbations. To do so, we performed transcriptome-wide differential expression relative to intergenic controls for each guide and correlated the results to the differential expression induced by other guides targeting the same gene^22^. This approach is comparable to prior assessments of loss-of-function screening modalities that examine the correlation of constructs that target the same gene^25^. In this Perturb-seq experiment, we observed that guides targeting the same gene are more correlated than guides that target different genes for both CRISPRko (Figure 3d) and CRISPRi (Figure 3e), with similar performance overall between the two technologies and the strongest effect at Day 14 (AUC=0.93, 0.94 for CRISPRko, CRISPRi). Upon repeating this experiment in the A549 lung cancer cell line (Day 7), we likewise observed clear agreement among guides targeting the same gene (AUC=0.89, 0.90 for CRISPRko, CRISPRi) (Supplementary Figure 1c).

We further interrogated the strength of on-target signals in this dataset by quantifying agreement in perturbation effects among guides that target the same gene but with different modalities (Figure 3f). Despite a distinct mechanism of action, CRISPRko and CRISPRi guides targeting the same gene exert remarkably congruous perturbation effects (AUC= 0.85). In sum, we conclude that both modalities in this experiment yielded satisfactory on-target activity since perturbation effects reproduce across independent guides both within and across modalities.

### CRISPRi shows greater off-target activity

While CRISPR guides typically target a unique 20 nucleotide sequence in the genome, Cas9 can tolerate certain mismatches and thus enable guide activity at unintended sites with partial complementarity^26–28^. Guide off-target activity has been observed to be more frequent with the use of dCas9 fused to transcriptional modulators, as guides with only partial complementary can nevertheless associate with DNA for biologically-meaningful lengths of time^29,30^. We sought to use the full transcriptomic readout of this experiment to detect such off-target events. Since we observe consistent on-target effects, transcriptomic changes elicited by only one guide targeting a gene likely reflect an off-target effect. Further, measuring the reproducibility of these guide-specific signatures distinguishes robust off-target effects from experimental noise^31^. This unbiased approach to detect promiscuous guides from Perturb-seq data complements a recently published strategy to validate activity at predicted off-target sites^32^.

To examine off-target activity in this manner, we obtained the median correlation of each guide to others targeting the same gene and plotted this value against the guide’s self-correlation across samples. We selected the Day 7 and Day 10 samples since transcriptional effects are unlikely to be complete by Day 4, and the Day 14 data are slightly distinct in structure (Supplementary Note 1). Consistent perturbation effects across samples effectively measure reproducibility since, in addition to deriving from distinct experimental time points, these samples represent separate transductions. Consequently, off-target activity is indicated when a guide demonstrates a cross-sample correlation that is substantially stronger than its correlation with other guides directed at the same target gene. We observed that CRISPRko guides targeting the same gene are generally just as correlated as individual guides across samples, suggesting minimal off-target activity. In contrast, numerous CRISPRi guides exhibit stronger correlation across samples than with other guides targeting the same gene (Figure 4a). We then obtain the difference between the cross-guide and cross-sample correlation for all guides and recognize positive outliers as promiscuous (Figure 4b).

**Figure 4.**
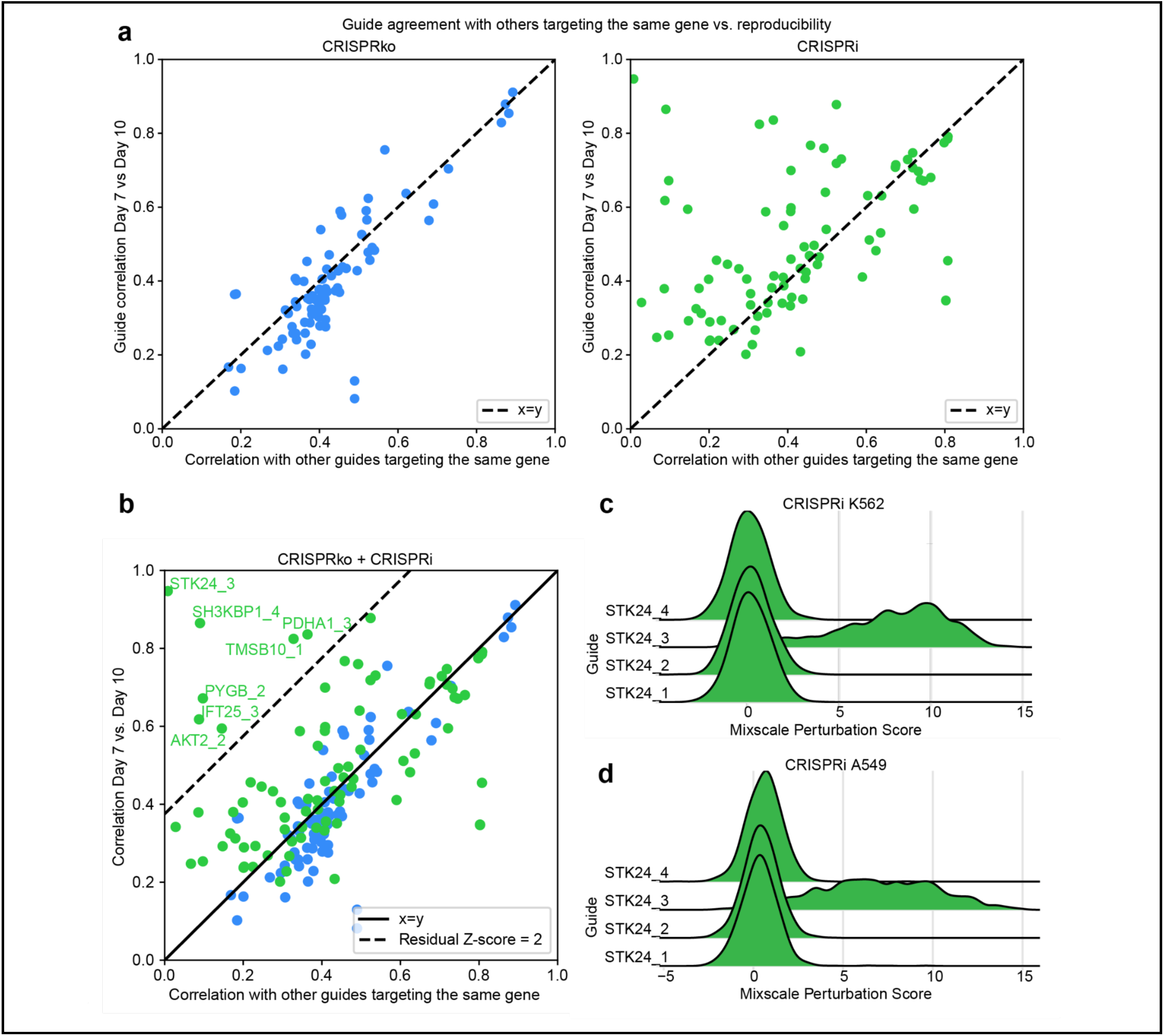
CRISPRi off-target activity observed in K562 Perturb-seq data. a) The Pearson correlation of each guide with other guides targeting the same gene with the same modality against the guide’s self-correlation across the samples collected at Day 7 and Day 10. The median cross-guide correlation is averaged across timepoints, and CRISPRko and CRISPRi guides are plotted separately. Correlation is calculated on signed p-values resulting from SCEPTRE. The dotted line represents the x=y line. b) Identification of promiscuous guides from the plots reported in a). Guides are z-scored relative to the distribution of all guide vertical residuals from the x=y line, and guides with a z-score above 2 are labeled in the plot. c) Receiver operating characteristic curve of correlation in signed p-values (SCEPTRE) between CRISPRko and CRISPRi guides that target the same vs. different genes at Day 14 in the K562 cell line. Correlation is calculated upon all transcripts that pass SCEPTRE pairwise crosswise correlation for both samples. The dotted line represents removal of CRISPRi guides STK24_3, SH3KBP1_4, and PYGB_2. d) Distribution of Mixscale perturbation scores in Day 7 K562 data among cells assigned to CRISPRi *STK24*-targeting guides. Cells are stratified by the assigned guide. e) d), in A549 data.

The guides we identify as promiscuous do not diverge from other guides targeting the same gene merely due to a lack of perturbation efficacy; rather, such guides mostly elicit a disproportionately high number of differentially expressed genes when screened in both the K562 (Supplementary Figure 2a) and A549 cell line (Supplementary Figure 2b). We additionally sought to validate that discordant guides reflect off-target instead of exceptional on-target activity. To do so, we reasoned that guide on-target activity promotes, whereas off-target activity deflates, the cross-modality agreement as presented in Figure 3f. Upon removal of guides we identify as promiscuous, the cross-modality agreement improves from AUC=0.85 to 0.87 (Supplementary Figure 2c). Thus, it is unlikely that the unique phenotypes elicited by these guides are related to target gene function.

Lastly, to confirm that our conclusions on guide promiscuity are not merely an artifact of our usage of SCEPTRE, we additionally analyzed this data with the Mixscale pipeline^33^. This approach models the inherent heterogeneity of knockdown efficacy across a cell population in order to boost statistical power during differential gene expression analysis. To accomplish this, Mixscale groups all cells carrying guides that target a certain gene and generates a vector from negative control cells. This vector represents the consensus effect of perturbing this gene. Next, for each individual cell in the group, a vector is generated from negative controls, and this vector is subsequently projected onto the gene vector to yield a scalar value, or “Mixscale score,” that indicates how well the cell agrees with the consensus effect. If all guides targeting a gene generate comparable (non-zero) Mixscale score distributions, this suggests that the guides agree on the transcriptomic effect of perturbation and thus successfully yield on-target activity. In contrast, if all cells with high Mixscale scores belong to an individual guide, that guide elicits a strong transcriptional program not shared with any other guides. We observed this phenomenon for CRISPRi *STK24* guide 3, thereby offering redundancy to conclusions drawn from off-target analysis with the results of SCEPTRE (Figure 4c). To assess if this guide behavior was context-specific, we also examined Mixscale score distributions in the A549 experiment and observed the same result (Figure 4d).

Ideally, CRISPRi guides with a propensity for off-target activity would be recognized and omitted *a priori*. The library presented here reflects avoidance of guides complementary to sequences proximal to the TSS of other genes. Studies have additionally attributed promiscuous dCas9 binding to enrichment of the GG motif in the 10-12 bases proximal to the PAM (otherwise known as the seed sequence)^3,32,34,35^. While this study features too few guides to meaningfully inform the relationship between guide composition and promiscuity, we note that guides with experimental evidence of off-target activity do not feature an excessive count of GG motifs in the seed sequence (Supplementary Figure 2d). Thus, strategic CRISPRi library design does not replace the need for guide-aware downstream analysis to avoid false positives.

### Both modalities risk false positives stemming from activity at the target site

Even after removal of promiscuous CRISPRi guides, the cross-modality agreement in Day 14 K562 perturbation effects (AUC=0.87) is not as strong as among guides that employ the same modality (AUC=0.93, 0.94 for CRISPRko, CRISPRi). For some genes, a decrease in transcript abundance and complete gene knockout could truthfully yield distinct phenotypes. We also consider that this observation reflects screening artifacts unique to either CRISPRko or CRISPRi. Unlike off-target activity inherent to the sequence of the guide, which can be mitigated by requiring agreement across guides targeting the same gene, here we focus on phenotypes that stem from activity at the intended site and are thus guide-agnostic.

CRISPRi neighbor knockdown has been observed for genes whose TSS is up to 1kb away from the TSS of a nearby gene, which comprises as many as 17% of protein-coding genes^3,9^. In theory, such genes cannot be assessed in isolation with CRISPRi since guides must target near the TSS in order to be effective^2^. In contrast, CRISPRko can target any exon of a gene, so the mere 3.3% of exons shared between genes can likely be avoided with deliberate library design^36^. Here, we identified genes targeted in our library with a neighboring TSS within a 1kb window and assessed expression of the neighboring genes relative to negative control cells. We observed that numerous CRISPRi guides reduce transcripts within 1 kb of their target site (SCEPTRE FDR <0.1)(Figure 5a). For example, in agreement with our viability screens in Figure 1c, we observed lower *NUDT21* transcript abundance in cells that received CRISPRi guides targeting the neighboring gene *OGFOD1*. No CRISPRko guides resulted in reduced expression of neighboring genes (Supplementary Figure 3), suggesting these observations were unrelated to loss of function of the target genes.

**Figure 5.**
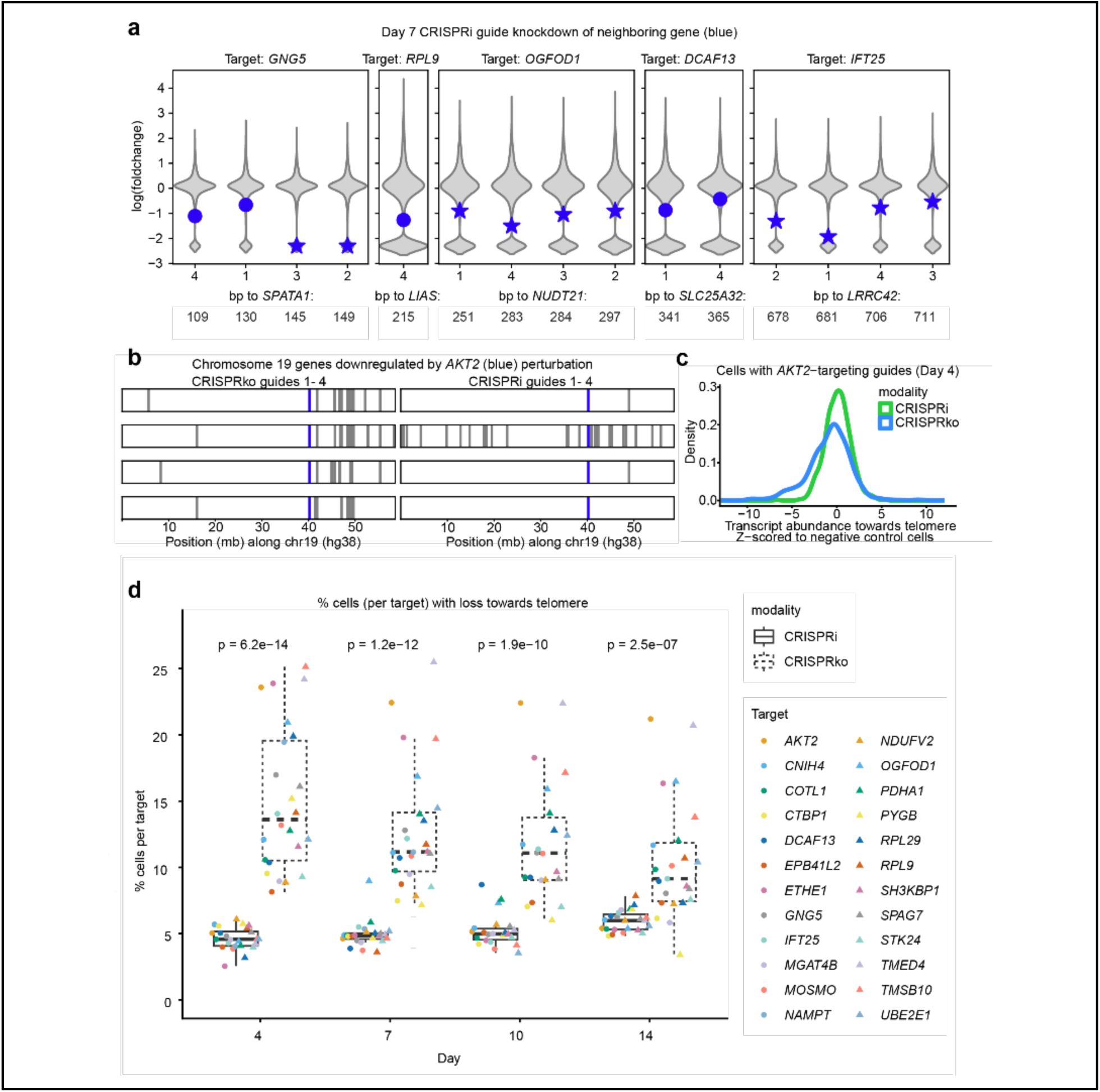
CRISPRko and CRISPRi feature distinct false positive modes stemming from activity at the target site. a) Transcriptional effect of CRISPRi guides targeting genes whose TSS is within 1kb of a neighboring gene TSS. Violin plots in grey indicate the Day 7 (K562) log fold change relative to intergenic-targeting guides of all transcripts that pass SCEPTRE pairwise quality control. Neighboring transcripts are shown in blue. Transcripts with significant knockdown (FDR <0.1) are denoted with a star marker. Guide distance to the neighboring gene MANE Select TSS is determined from the 17th position along the 20 nucleotide target site. b) Day 7 effect of *AKT2*-targeting guides along chromosome 19 in K562 cells. Genes marked in grey are significantly downregulated (SCEPTRE FDR<0.1, fold change <1) by any guide targeting the gene. The position of *AKT2* is shown in blue. c) Module score of genes towards the telomere from *AKT2* in cells with *AKT2* perturbed relative to cells with intergenic control guides. Module score is determined from transcriptomic counts at Day 4. Cells are stratified by modality. d) Per-sample fraction of cells with each target gene perturbed with putative chromosome arm loss. Loss is defined as lower detection of genes on the target chromosome arm towards the telomere than the bottom fifth percentile of negative control cells. For each time point, the distribution of loss rates from each modality are compared with a two-sided Mann-Whitney test and the resulting p-values are reported on the plot.

CRISPRko can also yield screening artifacts that cannot be revealed by examining agreement among guides targeting the same gene. In particular, CRISPRko may elicit effects unrelated to target function by introducing chromosomal instability. Prior studies have found that CRISPRko tends to induce downregulation of genes towards the telomere on the target gene chromosome arm in a manner that is not reproduced by CRISPRi^6^. Cells that have sustained chromosome-scale alterations by CRISPRko often exhibit a proliferative disadvantage, thereby potentially decreasing the harm of this artifact at later time points^7^. We first assessed this phenomenon in our data by examining the results of differential expression analysis for genes towards the telomere of library targets in perturbed cells. We noticed patterns consistent with DSBs driving chromosomal effects; for example, many transcripts downregulated by *AKT2-*targeting CRISPRko guides are on the q arm of chromosome 19 towards the telomere from *AKT2* (Figure 5b). CRISPRi guides targeting *AKT2* show few downregulated genes towards the telomere with the exception of guide 2, which elicits a disproportionate transcriptional effect consistent with its previously observed evidence of off-target activity (Figure 4b).

To identify the rate at which this aberrant DSB response occurs, we examined the transcript abundance of genes towards the telomere from the target in each cell. Returning to the example of *AKT2* perturbation, we observed loss towards the telomere in a subset of cells perturbed with CRISPRko but not CRISPRi (Figure 5c). Upon analysis of the entire dataset, we found that CRISPRko-perturbed cells show reduced expression of transcripts between the locus targeted by the guide and the telomere across all target genes and timepoints (Figure 5d). The frequency of this phenomenon peaks at Day 4, where an average of 15.0% of CRISPRko-perturbed cells exhibited lower expression compared to negative controls. The effect appears more modest by Day 14, which could be due to the proliferative disadvantage of cells with CRISPRko-induced chromosomal loss^7^. Nevertheless, this result provides further evidence that the cellular response to DNA double-stranded breaks with CRISPRko must be accounted for in subsequent phenotypic interpretation.

## Discussion

The high dimensionality of Perturb-seq screens presents an opportunity to compare insights on gene function gained with distinct loss-of-function techniques. However, published studies to-date have largely employed few constructs per gene, with some as low as one construct per gene, preventing the ability to identify legitimate perturbation effects via guide consistency. Here we present Cas9 CRISPRko and CRISPRi benchmarking libraries designed to enable direct comparison of these modalities. The libraries target 25 functionally diverse genes at four guides per gene, a design that enables the evaluation of guide concordance as a readout for perturbation efficacy. The libraries prioritize nonessential genes to support experimental readouts at late time points. Both modalities produced consistent, reproducible transcriptional signatures across guides and libraries, confirming robust on-target activity in the two cell lines evaluated. Further, cross-technology comparison showed substantial agreement between gene-level phenotypes.

CRISPRi guides more frequently exhibited off-target activity as measured by transcriptional signatures not shared with others targeting the same gene. False positives due to guide off-target activity are relatively straightforward to avoid in Perturb-seq experimental design and analysis; by incorporating numerous for each gene in a library, one can verify that perturbation effects reproduce across guide. This approach has been a cornerstone in the analysis of CRISPR screens with unidimensional readouts for well over a decade^37^. Notably, extra care is required when employing individual constructs that express two guides targeting the same gene^10,38,39^; while this may improve on-target knockdown, it also doubles the off-target liability of each reagent.

We particularly emphasize the importance of checking for guide agreement when implementing the Mixscale pipeline^33^, for an off-target phenotype can be interpreted as the on-target phenotype. For example, among cells with *STK24* knockdown by CRISPRi, only those carrying the guide that we have identified as promiscuous receive high Mixscale scores. If one were to proceed with weighted differential expression analysis using Mixscale scores, the promiscuous guide would disproportionately impact the result and likely result in misguided (pun intended) conclusions on gene function. It is therefore advisable to examine guide-level distributions of Mixscale scores when using this approach.

These modalities additionally diverge in failure modes that occur regardless of the guide selected. Such false positives require more specialized approaches to isolate genuine loss-of-function effects. In this study, a meaningful fraction of cells perturbed with CRISPRko demonstrated low detection of genes telomeric to the cut site, likely indicating large-scale chromosomal modifications. Such cells can be omitted from downstream analysis in the context of Perturb-seq screens. For CRISPRi, knockdown of genes with proximal TSSs presents challenges in the interpretation of phenotypes elicited by any guides targeting gene-dense loci. Since we observe relatively consistent neighbor knockdown, it is perhaps appropriate for analytical pipelines to assume that CRISPRi guides perturb all genes with TSS within 1kb of the target site.

One limitation of this study is that targets were selected based on prior evidence of transcriptional phenotypes in CRISPRi screens^10^, which may enrich for CRISPRi-associated features but underrepresent those associated with CRISPRko. Expanding the target space would enable a more comprehensive comparison of the modality-specific on-target performance. Additionally, some gene perturbations that induce modest transcriptomic effects may be better resolved by examining changes in chromatin accessibility^40^, image-based phenotyping^41^, or high-throughput protein assessment^42^.

As Perturb-seq data are increasingly implemented at atlas-scale and leveraged to develop foundation models of gene function and cell state, the artifacts described here represent a substantial risk to the interpretation of results. Reassuringly, the approaches presented here to mitigate such effects require relatively minimal computational effort and instead rely heavily on experimental design parameters. Moreover, given the described tradeoffs of each modality, we highlight that screening with both CRISPRko and CRISPRi when possible can elucidate robust loss-of-function phenotypes to confidently associate genes with their functions.

## Methods

### Library Design

Genes with high potential for strong perturbation effects are identified by analysis of existing genome-scale K562 Perturb-seq data at Day 8 post-transduction^10^. Genes are filtered to those well expressed in control cells (in the top 10% of transcripts) with p<0.001 in the reported energy distance test, which reports the significance of the transcriptome-wide effect size relative to negative control cells, and more than 50 differentially expressed genes (Anderson-Darling test, p<0.05). Gene essentially is determined by median Chronos scores across all cell lines featured in DepMap, and adequate expression is determined with gene transcript counts per million (DepMap 2026Q2).

Guides were selected with the CRISPick web tool in March 2024 with Human GRCh38 (Ensembl v.113) as the reference genome. Both libraries employ the same set of ten intergenic-targeting guides. Guide sequences are supplied with this manuscript (Supplementary Data 1).

### Mammalian cell lines and culture

K562 (female), A549 (male), and HEK293T (female) cells were obtained from the Cancer Cell Line Encyclopedia at the Broad Institute. All cells regularly tested negative for mycoplasma contamination and were maintained in the absence of antibiotics except during screens and lentivirus production, during which media was supplemented with 1% penicillin–streptomycin. Cells were passaged every 2–3 days to maintain exponential growth and were kept in a humidity-controlled 37 °C incubator with 5.0% CO2. A549 cells were passaged with DMEM supplemented with 10% Fetal Bovine Serum (FBS). K562 cells were passaged with RPMI supplemented with 10% FBS. HEK293T cells were passaged with DMEM supplemented with 10% heat-inactivated FBS.

### Modular vector design

Vectors were designed with the Fragmid web tool^17^. All pF plasmids were made by gene synthesis into the EcoRV site of pUC57-Kan (Genscript). All pF vectors were assembled into plasmids using Golden Gate cloning. pMV_AA088 was used to establish Cas9 for the CRISPRko experiment, and pRDB_332 was used to establish dCas9 for the CRISPRi experiment. Both guide libraries were cloned into the pRDB_422 CROP-seq vector.

pMV_AA088 (Addgene:216104): EF1α promoter expresses SpyoCas9 and blasticidin resistance; split by a 2A site (pRDA_722 destination vector (Addgene:216077); pF_AA179; pF_AA065; pF_AA012; pF_AA007; pF_AA009; pF_AA006).

pRDB_332: EF1α promoter expresses deactivated SpyoCas9, ZIM3 KRAB from Alerasool et al. 2020, and blasticidin resistance; split by a 2A site (pRDA_722 destination vector (Addgene:216077); pF_AA179; pF_AA065; pF_AA089; pF_AA032; pF_AA462; pF_AA006).

pRDB_422: CMV-driven CROP-seq vector, EF1α promoter expresses puromycin resistance^43^ (pRDA_789 destination vector (Addgene:216078); pF_AA065; pF_AA068; pF_AA175; pF_AA627).

### Modular vector destination vector pre-digest

The destination vector (pRDA_722) was pre-digested using BbsI and NEBuffer 2.0 (New England Biolabs) at 37°C for 2 h. The linearized destination vector was gel purified using 0.7% agarose gels and extracted with the Monarch DNA Gel Extraction Kit (New England Biolabs), before further purification by isopropanol precipitation.

### Modular vector Golden Gate assembly

The pF vectors were diluted to 10 nM in sterile water and cloned into a pre-digested destination vector via Golden Gate cloning. Each reaction contained 3 μL BbsI (New England Biolabs), 1.25 μL T4 ligase (New England Biolabs), 3 μL of 10x T4 ligase buffer (New England Biolabs), 75 ng destination vector, and a 1:1 molar ratio of fragments:destination vector. Reactions were carried out under the following thermocycler conditions: (1) 37°C for 5 min; (2) 16°C for 5 min; (3) go to (1), x100; (4) 37°C for 30 min; (5) 65°C for 20 min. The Golden Gate product was treated with Exonuclease V (New England Biolabs) at 37°C for 30 min before enzyme inactivation with the addition of EDTA to 11 mM. Per reaction, 10 μL of product was transformed into Stbl3 chemically competent E. coli (Invitrogen) via heat shock, and grown at 37°C for 16 h on agar with 100 μg/mL carbenicillin. Colonies were picked and grown at 37°C for 16 h in 5 mL Luria-Bertani (LB) broth with 100 μg/mL carbenicillin. Plasmid DNA (pDNA) was prepared (QIAprep Spin Miniprep Kit, Qiagen). Purified plasmids were verified by restriction enzyme digest and whole plasmid sequencing through Plasmidsaurus.

### Library production

Oligonucleotide pools of 110 guides were synthesized by Twist. BsmBI recognition sites were appended to each guides sequence along with the appropriate forward and reverse overhang sequences (bold italic) for cloning into the sgRNA expression plasmids, as well as primer sites to allow differential amplification of subsets from the same synthesis pool. The final oligonucleotide sequence structure was thus: 5′-[Forward Primer]CGTCTCA***CACC***G[guide, 20 nt]***GTTT***CGAGACG[Reverse Primer]-3’.

Primers were used to amplify individual subpools using 25 μL 2x NEBnext PCR master mix (New England Biolabs), 1-2 μL of oligonucleotide pool (∼30-300 ng), 5 μL of primer mix at a final concentration of 0.5 μM, and water to a final volume of 50 µL. PCR cycling conditions: (1) 98°C for 1 min; (2) 98°C for 30 sec; (3) 53°C for 30 s; (4) 72°C for 30 s; (5) go to step 2, 6-14x depending on library size; (6) 72°C for 5 min.

The resulting guide amplicons were PCR-purified (Qiagen). sgRNAs and iBars^43^ (Supplementary Data 1,2) were cloned into the pRDB_422 vector via three-way Golden Gate cloning with Esp3I (Fisher Scientific) and T7 ligase (Epizyme) under the following thermocycler conditions: (1) 37°C for 5 min; (2) 20°C for 5 min; (3) go to step 1, x99; (4) 37°C for 30 min; (5) 65°C for 10 min. The ligated product was isopropanol precipitated and electroporated into Stbl4 electrocompetent cells (Invitrogen) and grown at 37°C for 18 h on agar with 100 μg/mL carbenicillin. Colonies were scraped and plasmid DNA (pDNA) was prepared (HiSpeed Plasmid Maxi, Qiagen). To confirm library representation and distribution, the pDNA was sequenced by Illumina MiSeq.

### Lentivirus production

For small-scale virus production, the following procedure was used: 18 h before transfection, HEK293T cells were seeded in 6-well dishes at a density of 1×10^6^ cells per well in 1 mL of DMEM + 10% heat-inactivated FBS. Transfection was performed using the TransIT-LT1 transfection reagent (Mirus) according to the manufacturer’s protocol. Briefly, for each construct, 324μL of Opti-MEM (Corning) and 17μL LT1 was combined with a DNA mixture of the packaging plasmid pCMV_VSVG (125 ng; Addgene #8454), psPAX2 (1250 ng; Addgene #12260), and the transfer vector (312.5 ng). The solutions were incubated at room temperature for 30 min and added dropwise to cells. Plates were then transferred to a 37°C incubator for 6–8 h, after which the media was removed and replaced with DMEM + 10% FBS media supplemented with 1% BSA. Virus was harvested and filtered 40 hours after this media change.

A larger-scale procedure was used for pooled library production. 18 h before transfection, 18×10^6^ HEK293T cells were seeded in a 175 cm^2^ tissue culture flask and the transfection was performed the same as for small-scale production using 6 mL of Opti-MEM, 305 μL of LT1, and a DNA mixture of pCMV_VSVG (5 μg), psPAX2 (50 μg), and 10 μg of the transfer vector. Following addition of the transfection mix, flasks were transferred to a 37°C incubator for 6–8 h, then the media was aspirated and replaced with BSA-supplemented media; virus was harvested and filtered 40 h after this media change.

### Determination of lentiviral titer

To determine lentiviral titer for transductions, cell lines were transduced in 12-well plates with a range of virus volumes (e.g., 0, 150, 300, 500, and 800 μL virus) with either 1.5×10^6^ or 3×10^6^ cells per well in the presence of polybrene. For transduction, the plates were centrifuged at 821 x g for 2 h, after which 2 mL of warm media was added to reduce viral toxicity. Plates were then transferred to a 37°C incubator for 4–6 h. Each well was then trypsinized and pooled. Two days post-transduction, we seeded an equal number of cells into two wells of a 6-well plate, and added puromycin to one well. The following passage, both wells were counted for viability. A viral dose resulting in ∼30% transduction efficiency, corresponding to an MOI of ∼0.35, was used for subsequent library screening.

### Small molecule dosages

The dosages for the selection drugs blasticidin and puromycin were as follows for the relevant cell lines:

A549: blasticidin 5 μg/mL; puromycin 1.5 μg/mL
K562: blasticidin 8 μg/mL; puromycin 1 μg/mL

Blasticidin selection was completed over 12-14 days, and puromycin selection was completed over 5-7 days.

### Lentiviral transduction to establish stable cell lines

In order to establish stable Cas-expressing cell lines, K562 cells or A549 cells were transduced with a lentiviral library with pMV_AA088 (Cas9, for CRISPRko) or pRDB_332 (dCas9, for CRISPRi) in the presence of polybrene at a dosage of 1 µg/mL. Cells were centrifuged at 821 x g for 2 h in 12-well plates, after which 2 mL of warm media was added to reduce viral toxicity. Plates were then transferred to a 37°C incubator for 4–6 h. Replicate wells were then trypsinized and pooled. Successfully infected cells were selected for with blasticidin as described above.

### Pooled screens

For pooled screens, cells were transduced with a lentiviral library cloned into the pRDB_422 CROP-seq vector. Transductions were performed at a low multiplicity of infection (MOI ∼0.35), using enough cells to achieve a representation of at least 500 transduced cells per guide assuming a 20% −40% transduction efficiency. The transduction protocol was the same as listed above. Puromycin was added 1 day post-transduction. Cells were passaged on puromycin for 2 passages to ensure complete removal of non-transduced cells. After selection, cells were passaged at a minimum of 500x representation every 2-3 days. We repeated these transductions sequentially such that samples 4, 7, 10, and 14 days post-transduction were collected at the same time.

For the viability screen, cells were pelleted by centrifugation, resuspended in PBS, and frozen promptly for genomic DNA isolation.

### Genomic DNA isolation, PCR, and sequencing

Genomic DNA (gDNA) was isolated for the viability screen using either the KingFisher Flex Purification System with the Mag-Bind Blood & Tissue DNA HDQ Kit (Omega Bio-Tek), per the manufacturer’s instructions. The gDNA concentrations were measured by Qubit.

For PCR amplification, unless otherwise noted, gDNA was divided into 100 μL reactions such that each well had at most 2 μg of gDNA. Plasmid DNA (pDNA) was also included at a maximum of 100 pg per well. Each well of a 96-well PCR plate contained 1.5 μL of Titanium Taq (Takara), 10 μL of Titanium Taq buffer, 8 μL of dNTP Mix, 5 μL of DMSO, 0.5 μL of P5 primer at 100 μM stock, 10 μL of P7 primer at 5μM stock, and nuclease-free water added to 100 μL. PCR cycling conditions were as follows: (1) 95°C for 1 min; (2) 94°C for 30 s, (3) 52°C for 30 s, (4) 72°C for 30 s, (5) go to step 1, x31; (6) 72°C for 10 min. The forward primer bound to the vector in the U6 promoter with the sequence 5′-TTGTGGAAAGGACGAAACACCG-’3; the reverse primer bound to the vector in the trRNA with the sequence 5′-ACCGACTCGGTGCCACTTTTTCAA-’3. Each primer had additional sequences appended to the 5′ end necessary for Illumina sequencing. PCR products were pooled and purified with Agencourt AMPure XP SPRI beads according to manufacturer’s instructions (Beckman Coulter, A63880). Samples were sequenced using Illumina technology NovaSeq 6000 S Prime 100 cycles, single-read, with a 5% spike-in of PhiX. Guide sequences were extracted from sequencing reads by running PoolQ with the search prefix “CACCG”, and reads were counted by alignment to a reference file of all possible guides present in the library.

### Analysis of viability screen data

Following deconvolution, the resulting matrix of read counts was normalized to reads per million by the following formula: read per guide/total reads per condition×10^6^. Normalized counts were log2-transformed, and the Day 7 and Day 14 (post-transduction) values were subtracted from the plasmid DNA (pDNA) to obtain the log2 fold change for each guide.

### Single cell RNA-seq sample preparation and sequencing

The particle-templated instant partition sequencing (PIPseq) method was employed for single cell RNA sequencing. Two PIPseq cell encapsulations occurred on Day 4, 7, 10, and 14 post-transduction for K562 cells and Day 7 for A549 cells. Days 4, 7, and 10 featured a mixture of cells from the CRISPRko and CRISPRi experiments, each with 2 technical replicate PIP encapsulations (i.e. the same exact cell mixture went into two different PIP tubes). At Day 14 in K562 cells, there were discrete PIP encapsulations for the CRISPRko and CRISPRi experiments in place of technical replicates. Cells were encapsulated and lysed at the Broad Institute per the Illumina Single Cell 3′ RNA Prep T100 protocol.

The PIPs were then shipped at room temperature to Broad Clinical labs for downstream processing within 72 hours of encapsulation. Downstream steps were performed as written with 8 cycles of GEX PCR, 12 cycles for CRISPR WTA, and 10 cycles for CRISPR PCR. Samples were quantified using qPCR and Tapestation, then normalized and pooled at a ratio of 90% GEX to 10% CRISPR per PIP.

Pooled samples were sequenced across a lane of a Novaseq X 25B flowcell, 100 cycle kit with a read structure of 45×10x10×72.

### Alignment of sequencing data

The samples were processed with the CRISPR scRNA pipeline of Illumina DRAGEN software v4.4.5 with the following usage:

~~~
/opt/edico/bin/dragen --annotation-file /scratch/hg38-alt_masked/genes.gtf.gz --autodetect-
reference-validate true --dump-map-align-registers true --enable-cnv false --enable-map-align
true --enable-map-align-output false --enable-metrics-json true --enable-rna true --enable-
single-cell-rna true --enable-sv false --enable-variant-caller false --fastq-list
paired_fastq_list.csv --fastq-list-sample-id AC1 --force --generate-sa-tags true --ignore-input-
mate read1 --intermediate-results-dir /scratch --json-dataset-type
/opt/edico/config/datasettype.json --lic-instance-id-location /opt/instance-identity --logging-
to-output-dir true --output-directory SAMPLENAME --output-file-prefix SAMPLENAME --qc-enable-
depth-metrics false --ref-dir /scratch/hg38-alt_masked/DRAGEN/11 --repeat-genotype-enable false -
-rna-library-type SF --rrna-filter-enable true --scrna-barcode-position 0_7+11_16+20_25+31_38 --
scrna-barcode-sequence-list ealr_barcode_whitelist.txt --scrna-enable-pipseq-crispr-mode true --
scrna-enable-pipseq-mode true --scrna-feature-barcode-groups CACTCAATTC.CACAACTTAA.1 --scrna-
feature-barcode-reference FEATURE-BARCODE-REFERENCE-FILE --scrna-umi-position 39_41 --single-
cell-threshold inflection --umi-source qname --validate-pangenome-reference false --vc-enable-
profile-stats true
~~~

The resulting analysis is conducted only on cells deemed as “passing” by DRAGEN, which excludes those with disproportionately low UMI or gene counts as these are understood to not represent an entire cell. DRAGEN summary files are used to inform the per-sample cell count, median genes per passing cell, and median transcriptomic UMIs per passing cell reported in Table 1 of this manuscript.

### Filtering to cells with sufficient quality

As previously mentioned, we omitted data from the K562 Day 4 technical replicate 1 sample due to low detection of transcriptomic reads. We believe this discrepancy to be a result of error in sample processing or library preparation instead of sequencing; whereas the total number of input reads to alignment was comparable to that of other samples, only 12.4% were classified by DRAGEN as barcoded gene expression reads.

Using the Seurat R package, we retain genes expressed in at least three cells per sample. We concatenated the set of cells from all remaining technical replicates. Cells for which at least 15% of transcripts were mitochondrial were excluded since these may represent dying cells.

### Guide Assignment

CROP-seq vectors capture the guide sequence in the RNA Pol II transcript, thereby enabling guide detection with the addition of the guide library to the reference transcriptome prior to alignment. However, this experiment leverages the CRISPR Direct Capture feature counts to determine guide assignments. The following approach was applied for each sample:

1. Remove cells with guide purity below 65%. Guide purity is calculated here as the number of feature counts belonging to the most abundant guide in the cell divided by the total count of guide feature counts in the cell. Thus, a guide purity of 65% indicates that 65% of guide feature counts in a cell belong to a single guide. This step ideally removes instances of doublet encapsulation and droplets that contain only ambient guides.
2. Remove cells with 0 guides that have enough feature counts to represent a native guide. To determine the minimum number of feature counts for a native cell, we recommend leveraging elbow plots in which the x-axis is a feature count and the y-axis is the number of guide:cell pairings below that count. Since each cell should only feature a single native guide and thus the majority of all possible guide:cell pairings in the experiment are not a true assignment (merely reflecting ambient guides), the ideal threshold is the “elbow” of the plot: the lowest feature count that excludes the majority of pairs. For the experiments described in this manuscript, this threshold ranges from 10-14 feature counts.
3. Among the cells remaining, assign the most abundant guide.
4. Remove guides (and, consequently, cells carrying such guides) assigned to fewer than 100 cells since guides with inadequate coverage demonstrate suboptimal performance in capturing perturbation signatures ^21^.

This approach yields assignment rates ranging from 72.69% to 90.02% across samples for the experiments described in this manuscript (Table 1).

### Differential expression analysis

Differential expression analysis was performed separately for each sample to inform guide perturbation effects. SCEPTRE low MOI ^22^ was run in singleton mode on raw sequencing counts to identify genes differentially expressed in cells with each guide relative to cells with intergenic-targeting negative control guides. This method was employed because it uses a permutation-based approach to avoid model misspecification, a relevant concern given the sparsity of single cell transcriptomic data. The n_nonzero_trt_thresh argument of the run_qc method was reduced from its default value of 7 to 0 to enable detection of transcripts with complete loss in perturbed cells. Otherwise, all default pipeline settings were employed. For downstream analysis, the fold change reported from SCEPTRE is incremented prior to manually calculating log10(fold change +0.1) to prevent division by 0 for transcripts absent in perturbed cells.

### Quantification of guide similarity

For each guide, the p-value of each transcript that passed quality control from SCEPTRE is log-transformed and signed according to its fold change. For every pair of guides, Pearson’s correlation is calculated on the intersection of transcripts that passed quality control for both guides.

### Identification of promiscuous guides

Guide promiscuity is initially determined using the samples collected at Day 7 and 10 post-transduction in the K562 cell line. For each guide, the signed p-values derived from SCEPTRE are correlated (Pearson’s) with other guides targeting the same gene with the same modality from the same sample. These cross-guide correlations are collapsed with the median as to avoid penalizing non-promiscuous guides that share a target with a promiscuous guide. These median correlations are then averaged across the Day 7 and Day 10 sample. This value is then subtracted from the guide’s self-correlation, namely, the correlation in signed p-values of the guide across the Day 7 and Day 10 samples. These differences are then z-normalized collectively for the CRISPRko and CRISPRi experiment, and guides with a z-score above 2 are identified as promiscuous.

### Mixscale

Mixscale^33^ is employed using the transcriptome of each cell reduced to ten principal components. The protocol identifies the 20 most similar negative control cells carrying intergenic-targeting guides in the generation of gene vectors.

### Quantifying CRISPRko arm loss

Chromosome arm positions were determined using the hg38 UCSC Chromosome Bands Localized by FISH Mapping Clones track^44^. Gene positions were determined from Ensembl v.115 annotations. Cells with *MPG* perturbed were excluded from this analysis since there are only three genes closer to the telomere. For the remainder of targets, the minimum number of genes in any condition after filtering for genes with sufficient coverage is 17.

Seurat’s AddModuleScore function was employed to identify the enrichment of genes towards the telomere on the target gene chromosome arm in each cell. Thus, the set of genes in this module depends on the gene targeted in the cell. To calculate enrichment at each time point, all cells in the sample are used as the reference set. Cells are determined to have evidence of loss of genes towards the telomere if the enrichment of towards-telomere genes is below the bottom fifth-percentile of negative control cells. For this analysis, negative controls are filtered to cells carrying intergenic guides 1,2,5, and 9, for these guides do not target the same chromosome as any of the library targets.

### Data visualization

Figures were created with the Python packages matplotlib, seaborn, adjustText and R packages ggplot, Seurat, and Mixscale. For all boxplots shown, boxes depict 25th (Q1) and 75th (Q3) percentiles as minima and maxima and the center represents the median; whiskers identify outlier points by depicting Q1 −1.5∗IQR and Q3 + 1.5IQR, where IQR represents the range between Q1 and Q3. Schematics were created with https://BioRender.com.

## Data and Code Availability

Aligned, quality-filtered Perturb-seq datasets with guide assignments have been deposited as RDS files to Zenodo at https://doi.org/10.5281/zenodo.20722064^45^. The results of differential expression analysis (SCEPTRE) per guide for all samples have been deposited to Zenodo at https://doi.org/10.5281/zenodo.21136747^46^. All original code and as well as the unprocessed data from the viability screen have been deposited to GitHub at github.com/ldrepano/head-to-head-CRISPRko-CRISPRi-Perturbseq and Zenodo at https://doi.org/10.5281/zenodo.21137515^47^.

## Acknowledgements

We thank Thouis R Jones, Evelyn Tong, Isabella Boyle, Orr Ashenberg, Elisa Donnard, and Aziz Al’Khafaji for guidance and feedback in the analysis of Perturb-seq data; Robert W Shea, Alia Yapo, Meghan Biggins, and Toni M Delorey for experimental assistance; Eleanor Kaplan for support in vector design; Ian Smith for guidance in methodology to recognize off-target effects in genetic screening data. This work was supported in part by the Functional Genomics Consortium.

## Author Contributions

Conceptualization: LMD, JGD

Investigation: LMD, BEV, AJC, DD, TS, JGD

Data curation: LMD, BEV, AJC, MG

Formal analysis: LMD, MG

Resources: HDR, DD, AWN, TS, PSW, JGD

Software: LMD, MG, AWN

Visualization: LMD, KBY

Supervision: HDR, DD, PSW, JGD

Writing (original draft): LMD

Writing (review & editing): LMD, SS, KBY, JGD

All authors read and approved the final manuscript.

## Supplementary Information

Supplementary Figures 1, 2, 3

Supplementary Note 1: Comments on shared intergenic controls across libraries.

Supplementary Data 1 [CRISPRko-CRISPRi-perturbseq-benchmark-guides.csv]: Guide sequences in the Perturb-seq library with their corresponding targets and indices for comparison to guide labels in the manuscript figures.

Supplementary Data 2[cas9_UMIRef_P6.csv]: Library of iBar UMIs cloned into the vectors screened in this library. Since this screen detects guides with Direct Capture technology instead of the Pol II transcript, the iBars are not recovered in this experiment and thus not utilized.

**Supplementary Figure 1, related to Figure 3.**
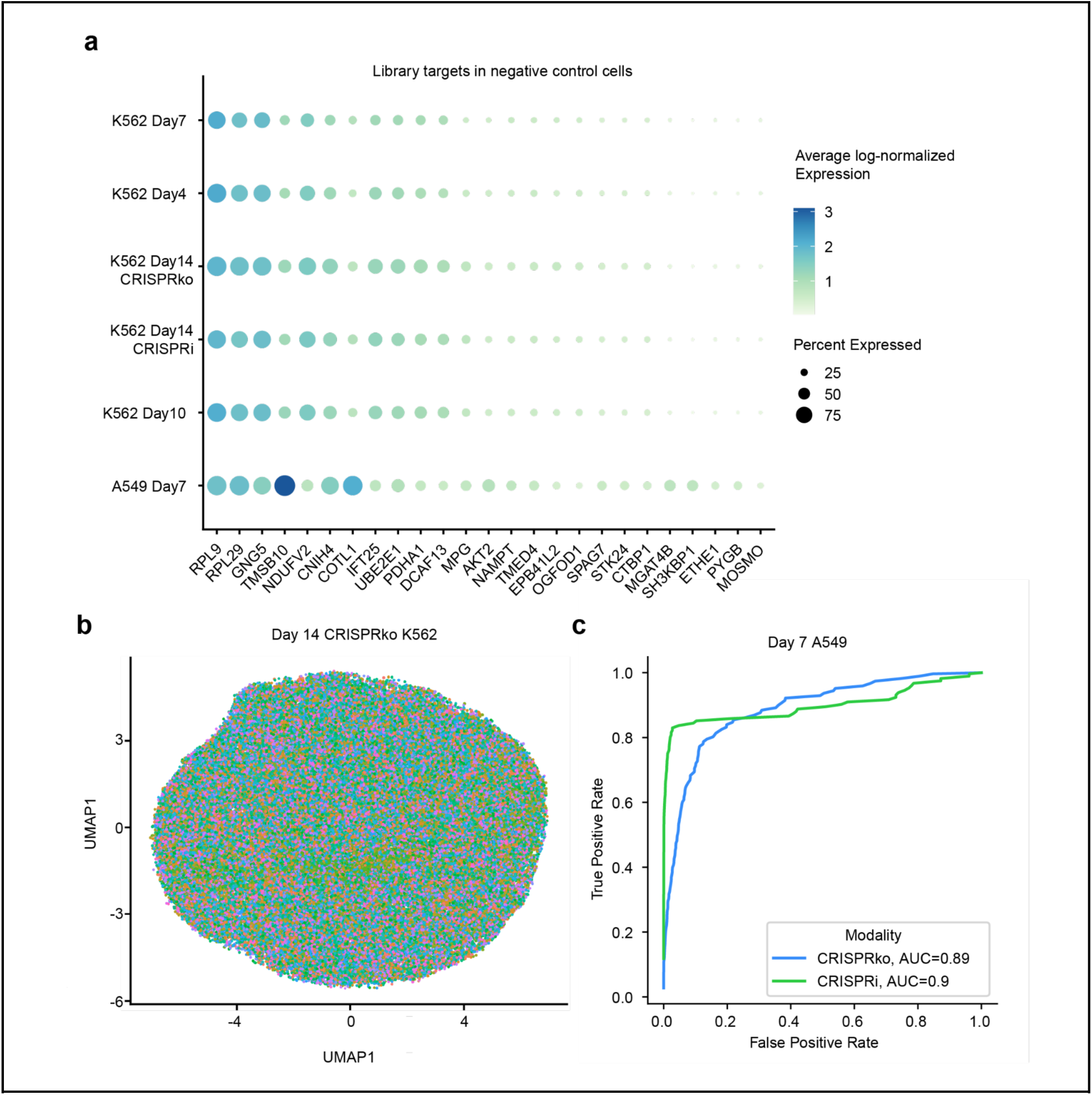
a) Average log-normalized counts of library target transcripts in cells carrying intergenic-targeting guides. b) UMAP analysis of all individual cells in the Day 14 CRISPRko K562 Perturb-seq sample. This analysis is conducted with the 2000 most highly variable genes and the use of 10 principal components and 8 neighbors. Cells are colored according to the target gene of the guide assigned. c) Receiver operating characteristic curve of correlation in signed p-values (as determined by SCEPTRE) for CRISPRko guides targeting the same vs. different genes in the A549 cell line at Day 7 post-transduction.

**Supplementary Figure 2, related to Figure 4:**
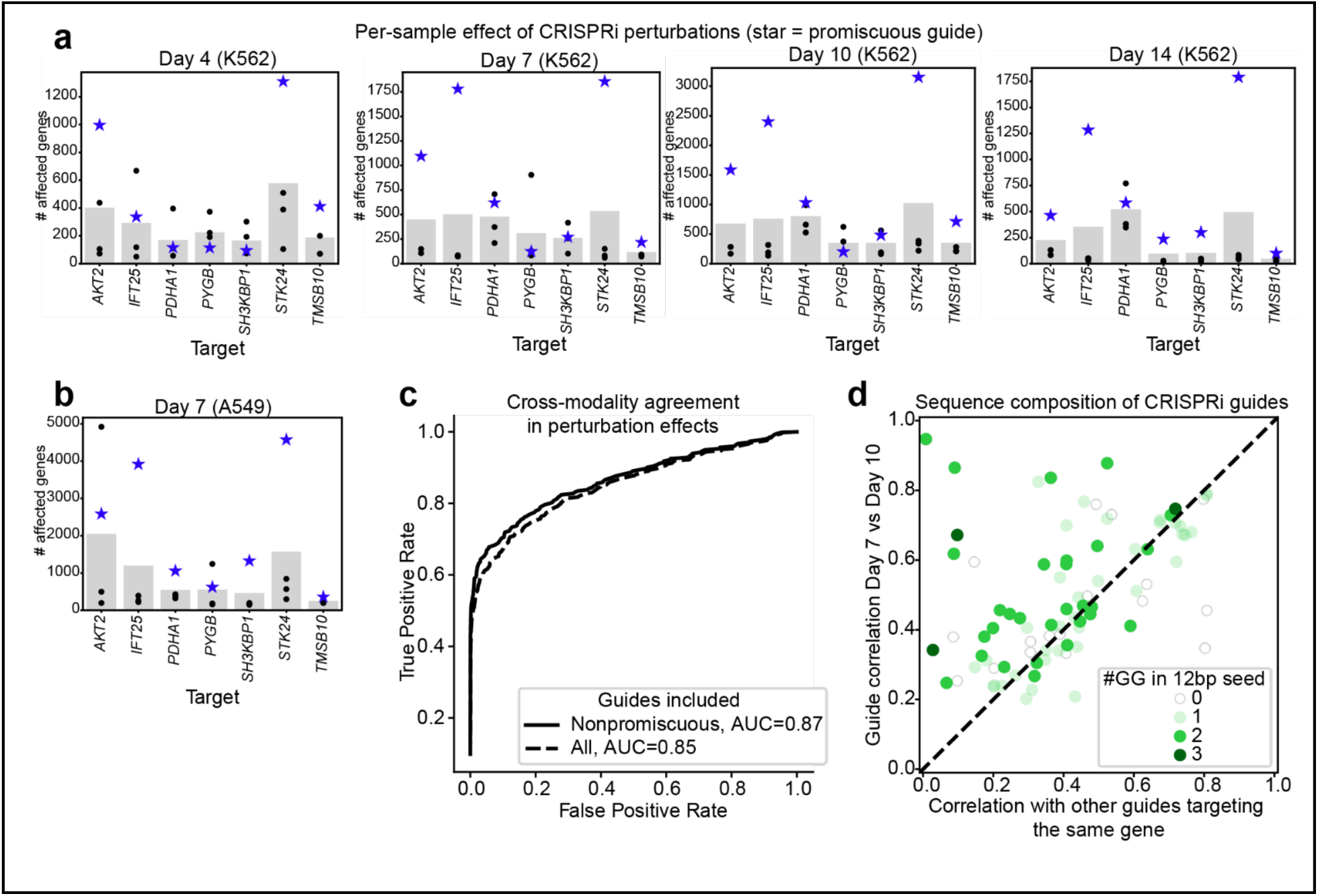
Characterization of CRISPRi guides identified as promiscuous. a) Count of differentially expressed response genes (SCEPTRE FDR <0.1) per guide, subset to targets of promiscuous CRISPRi guides. The bar height indicates the mean count of differentially expressed genes per target, and each point represents an individual guide. Guides identified as promiscuous in Figure 4b are denoted with a blue star marker. Each plot represents a distinct sample with a separate transduction to demonstrate reproducibility. The results represent data from the K562 cell line. b) a), but for the data collected at Day 7 in the A549 cell line. c) Figure 3f with the analysis repeated upon the removal of guides identified as promiscuous. d) Figure 5a, but with CRISPRi guides shaded according to the count of GG motifs in the 12 base pair seed sequence.

**Supplementary Figure 3, related to Figure 5a:**
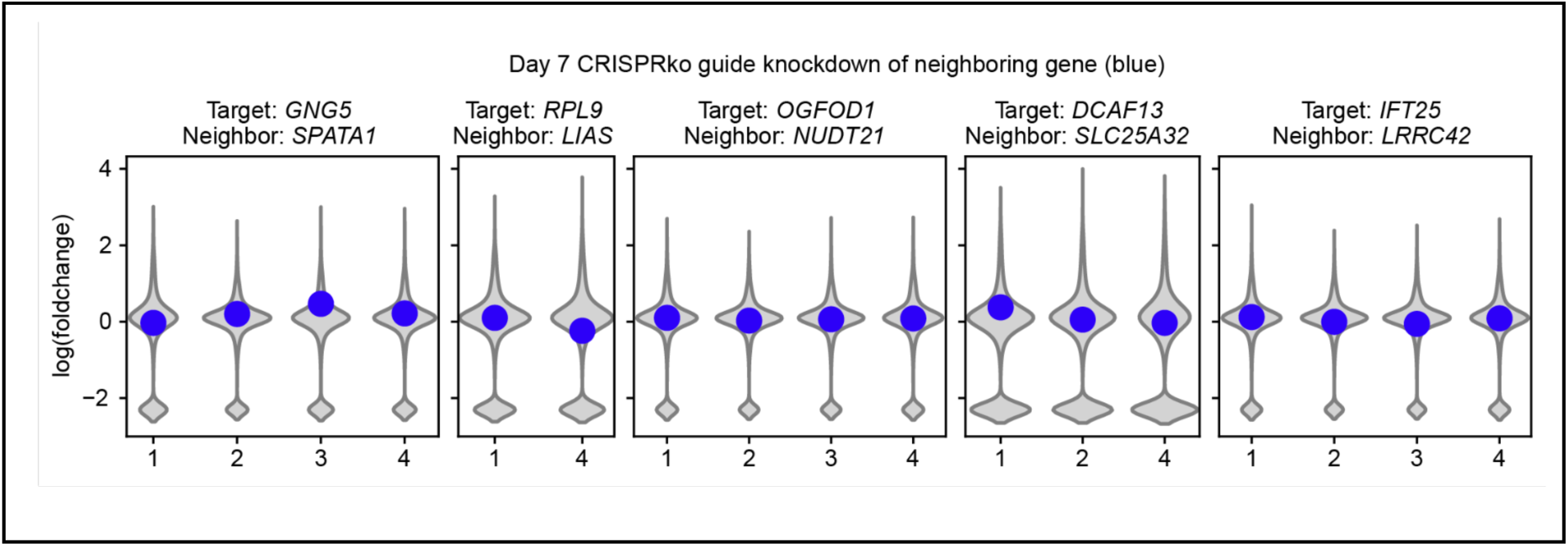
Transcriptional effect of CRISPRko guides targeting genes whose TSS is within 1kb of a neighboring gene TSS. Violin plots in grey indicate the Day 7 (K562) log fold change relative to intergenic-targeting guides of all transcripts that pass SCEPTRE pairwise quality control. Neighboring transcripts are shown in blue. Transcripts with significant knockdown (FDR <0.1) are denoted with a star marker.

**Supplementary Note 1**

Since both the CRISPRko and CRISPRi experiments utilize the same set of intergenic-targeting guides as negative controls, the perturbation effect of each gene-targeting guide is determined relative to a mixture of CRISPRko- and CRISPRi-perturbed cells (with the exception of the Day 14, for which we sequenced the experiments separately and thus can distinguish negative controls). Transcripts that are influenced by the presence of CRISPRko or CRISPRi consistently emerge as hits in the results of SCEPTRE differential expression analysis; for example, genes that score as upregulated for numerous targets in the CRISPRko experiment are also downregulated for many targets with CRISPRi. The opposite is true for genes with higher expression in cells perturbed with CRISPRi. These genes with apparent modality-specific expression are distinct for the K562 and A549 experiment.

Since our library did not include non-targeting guides, it is unclear if one or both of these modalities yields a transcriptional change from unperturbed cells. Nevertheless, one can mitigate this artifact when designing a Perturb-seq screen by simply employing negative control cells of the same modality.

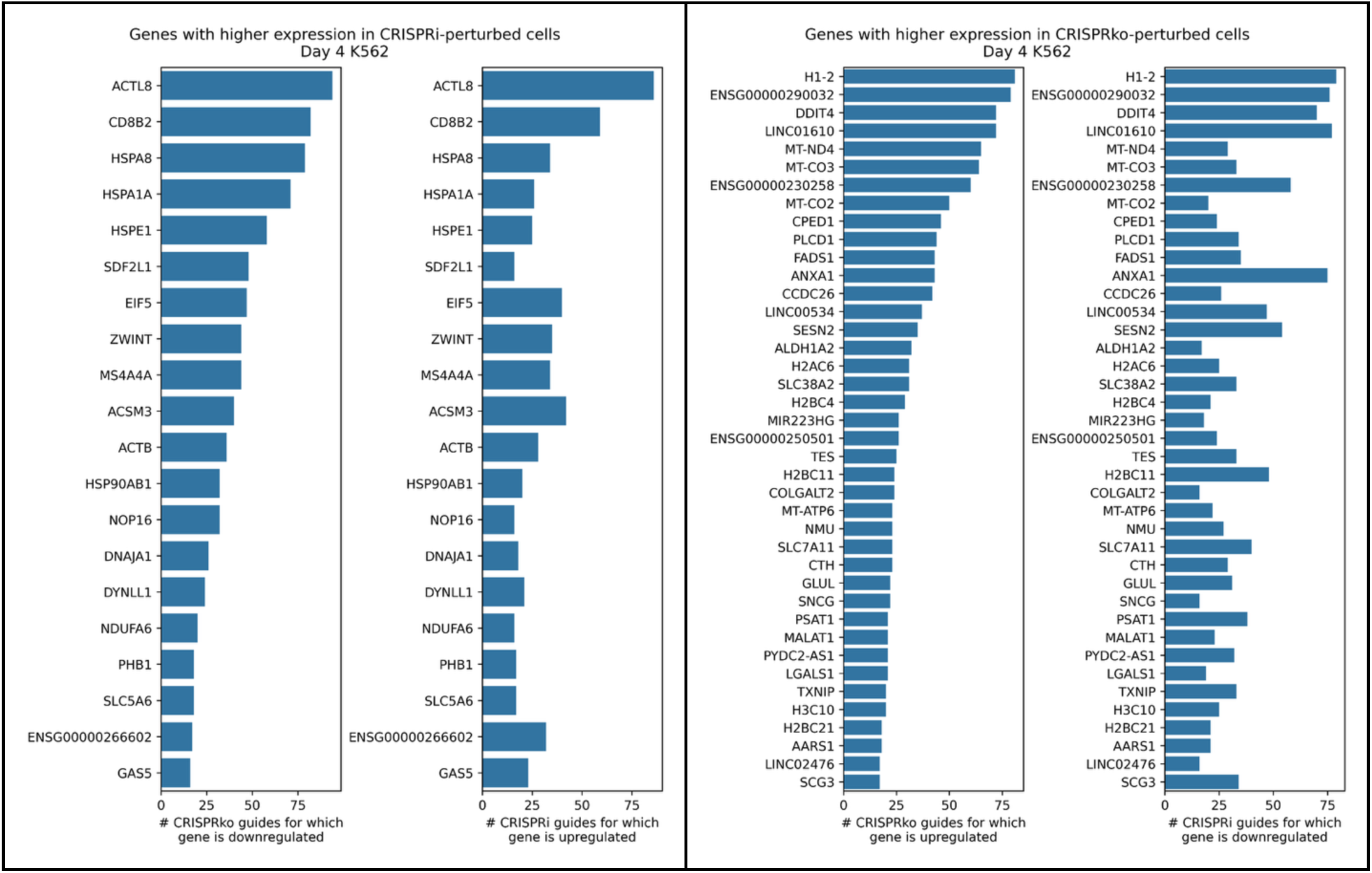
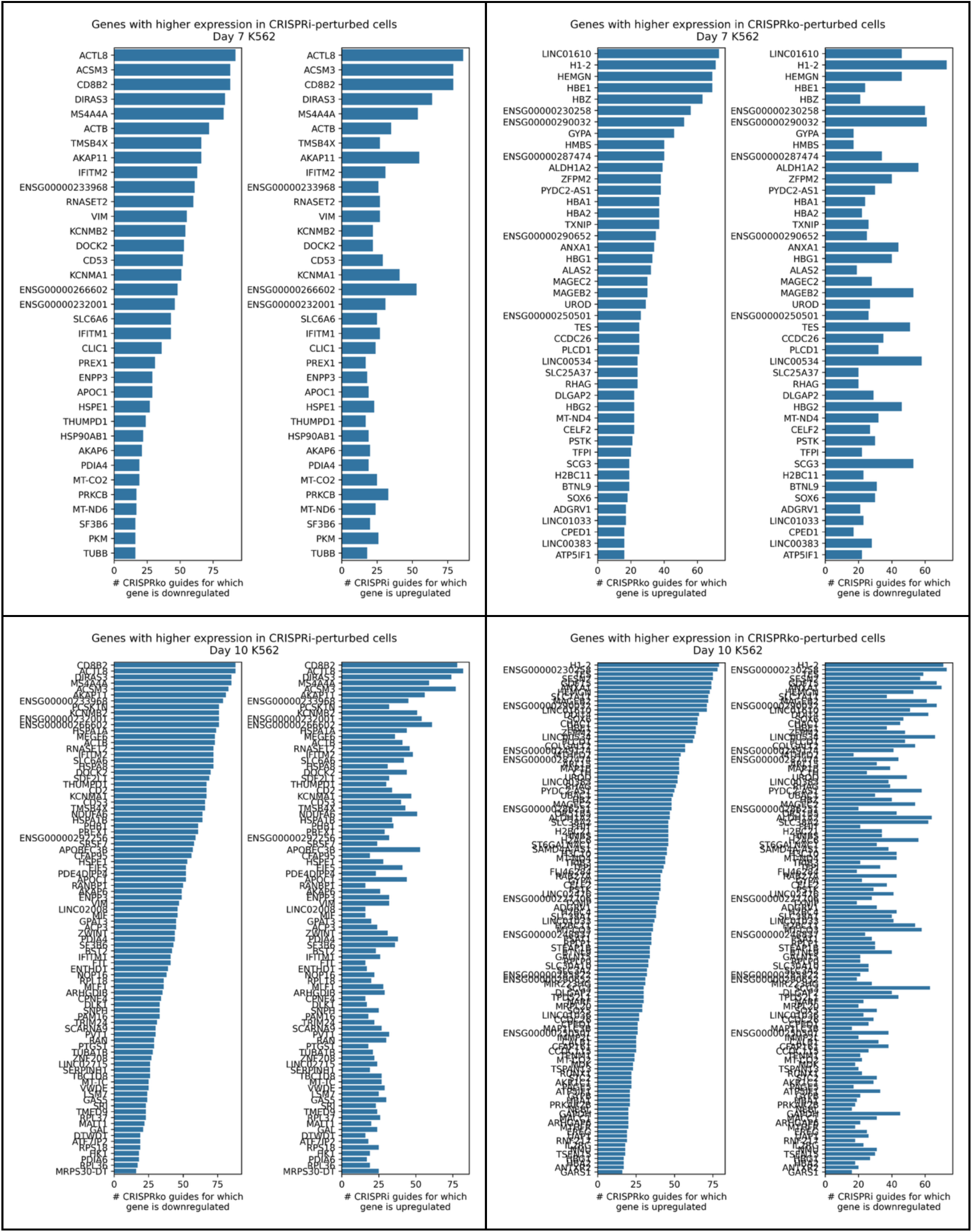
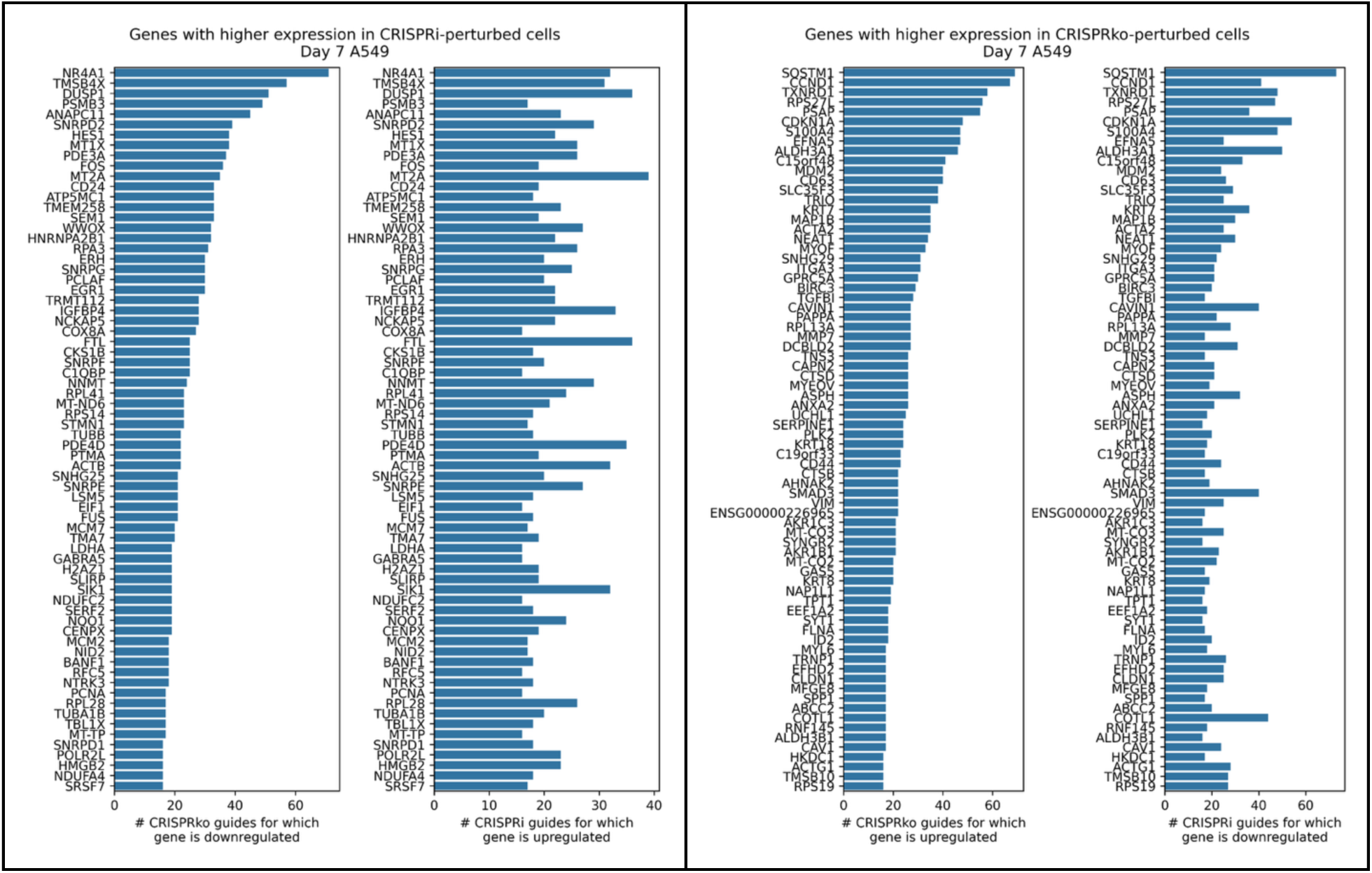
Differentially expressed genes are defined as those with SCEPTRE FDR <0.1. Genes are determined to be upregulated or downregulated depending if their fold change relative to intergenics is above or below 1. Only response genes differentially expressed in a particular direction for at least 15 guides per modality and sample are shown in the plots.

